# Bidirectional links between sleep regulation and cardiac physiology revealed by *Drosophila* modeling of genes at insomnia and cardiovascular disease loci

**DOI:** 10.1101/2025.04.07.647668

**Authors:** Torrey R. Mandigo, Farah Abou Daya, Dev Patel, Suraj Math, Lily Ober, Matthew Maher, Girish C. Melkani, James A. Walker, Richa Saxena

**Author notes:** These authors contributed equally to this work (Torrey R. Mandigo, Farah Abou Daya).

## Abstract

Insomnia is a common sleep disorder associated with negative long-term health outcomes, including cardiovascular disease (CVD). We selected 16 genes from 13 insomnia- and CVD-associated genetic loci and disrupted *Drosophila melanogaster* orthologs in neuronal or cardiac tissue to characterize their roles in regulating sleep and cardiac physiology. Neuronal disruption of four orthologs (*APOB*, *FUR*, *CYP17A1*, and *TCF4)* resulted in short-sleeping flies, and three (*MRAS*, *HDAC9*, and *TDRKH*) resulted in long-sleeping flies. Short-sleeping fly lines impacted cardiac physiology consistent with links between short or poor-quality sleep and increased CVD in humans. Conversely, heart-specific disruption of five orthologs (*RASD1*, *PHACTR1*, *CNNM2*, *MRAS*, and *TCF4)* led to defects in cardiac physiology, with varied effects on sleep. Heart-specific knockdown lines that altered fractional shortening, a measure of cardiac contractility, also influenced long and short sleep states. These findings reveal bidirectional relationships between sleep states and cardiac performance, representing a potential feedback loop linking insomnia and CVD.

**Teaser:** Disruption of insomnia and CVD-associated gene orthologs in *Drosophila* neurons and heart lead to alterations in sleep states and cardiac contractility that mutually reinforce poor sleep and poor cardiac performance.

## INTRODUCTION

Insomnia disorder, defined by persistent difficulty in initiating or maintaining sleep, and corresponding daytime dysfunction, occurs in 10-20% of the population, and leads to high socioeconomic costs and substantial lifetime morbidity (*1*). Various risk factors contribute to the development of insomnia, including irregular sleep schedules, poor sleep hygiene, stress, underlying medical conditions and genetic predisposition (*1*). The co-occurrence of insomnia with objective short sleep duration represents the most biologically severe phenotype of insomnia disorder (*2*). Longitudinal studies suggest that insomnia increases the risk for developing anxiety disorders, alcohol abuse, major depression, and cardio-metabolic disease (*3*). Cognitive-behavioral therapies are the recommended first-line treatments (*4*). In addition, pharmacologic treatments target synaptic neurotransmission (via GABAergic pathways), cortical arousal (via histamine receptors), or the melatonin system, but these drugs have variable effectiveness, may be habit-forming, and have debilitating side effects (*5*). A better understanding of the etiology and pathophysiological processes would enable identification of new personalized therapeutic strategies for insomnia, that may in turn impact downstream disorders. Family-based heritability estimates suggest that insomnia has a genetic component (22%–25%) (*6*) and novel genetic loci have recently been identified (*7–10*) but the effector genes at these genomic regions and related biological mechanisms that drive insomnia are poorly understood.

Multiple lines of evidence suggest that insomnia is an important risk factor for incident cardiovascular disease (*11*), the most significant contributor to mortality. Notably, between 2017-2021, 127.9 million US adults (48.6%) had some form of cardiovascular disease, and the American Heart Association has recently recognized sleep problems as one of eight essential lifestyle risk factors (*12*). Consistent association of insomnia with cardiovascular disease has been observed in cross-sectional and prospective epidemiologic studies (*3*, *13–15*). Insomnia with polysomnographic or self-reported short sleep duration is associated with incident cardiovascular disease (n=4,994, HR, 1.29 and HR,1.28) in the Sleep Heart Health Study (*16*), and similar findings for insomnia with actigraphy-measured or self-reported short sleep duration are seen in the UK Biobank (n=6,959, hazard ratio [HR]=1.39, 95% CI=1.11-1.74; (*2*). Genetic studies using Mendelian Randomization support a causal relationship, suggesting insomnia confers a 2-fold increased risk of coronary artery disease (*17*). Physiologic and experimental studies suggest that insomnia with objective short sleep duration may be associated with activation of the hypothalamic-pituitary-adrenal axis and increased neurocognitive-physiologic arousal, which may link insomnia to cardiovascular disease (*15*, *16*). However, it is currently unknown to what extent the increased CVD risk arises from shared underlying biological and genetic pathways, or whether the insomnia state leads to physiologic changes that ultimately result in CVD. Recent studies describe a mechanistic role of sleep in influencing atherosclerosis through hematopoiesis in mice (*18*) and link immune function to sleep in *Drosophila* (*19*), providing an important research framework to probe underlying mechanisms using model organisms.

Furthermore, epidemiologic and experimental studies suggest that the relationship between insomnia and cardiovascular disease is likely bidirectional. Insomnia frequently coexists with cardiovascular disease, and shared mechanisms may drive both phenotypes (*20*). For individuals with a history of cardiovascular events, those with insomnia experience higher rates of cardiac death and worsening heart failure than those without insomnia (*20*, *21*). Notably, myocardial infarction has been found to augment sleep through the recruitment of monocytes to the brain after cardiac injury in both mice and humans. This increased sleep was found to provide a protective effect against further cardiac damage as sleep disruption after myocardial infarction worsened cardiac outcomes (*22*). Further studies are required to fully elucidate the mechanisms linking sleep and cardiac function.

Model organisms, such as *Drosophila melanogaster*, provide the powerful genetic tools necessary to study sleep disruption and related cardiovascular function, and to identify and further probe candidate causal genes at genome-wide association studies (GWAS) loci for related insomnia and cardiovascular disease phenotypes. The fruit fly has been an excellent model system for basic discoveries concerning sleep and circadian rhythms and cardiovascular disease. A sleep-like state in *Drosophila melanogaster* comprises both homeostatic and circadian components (*23*, *24*). Mechanisms and regulation of sleep are deeply conserved from flies to humans (*25*, *26*). Recently, multiple studies have utilized *Drosophila* to elucidate the mechanisms underlying sleep GWAS signals (*27–29*). Furthermore, several quantitative measures of cardiovascular function in *Drosophila* have also been used to model cardiac disease (*30–32*). While the morphology of the *Drosophila* heart differs from that of mammals, many conserved pathways govern *Drosophila* and mammalian cardiac development and function (*30*, *32–34*). Furthermore, disruptions of orthologous genes associated with a higher risk of CVD in humans result in similar outcomes in flies (*35*). Notably, links between sleep and cardiac function are also beginning to be investigated in *Drosophila*. In a recent study, altered sleep and cardiovascular function were both found to increase levels of the *Drosophila* cytokine Upd3. Furthermore, overexpression of Upd3 in the brain was sufficient to alter cardiac function, while overexpression of Upd3 in the heart was sufficient to cause sleep fragmentation (*29*). Together, this research suggests that *Drosophila* is a well-suited model organism for investigating the link between insomnia and CVD.

Here, we prioritized a short list of human genes associated with insomnia and cardiovascular disease by querying genome-wide associations for insomnia, sleep disturbances and cardiovascular traits. We model disruption of Drosophila melanogaster orthologs in neurons and cardiac tissue to characterize their roles in regulating sleep and cardiac physiology.

## RESULTS

### Prioritization of insomnia- and CVD-associated candidate genes for *Drosophila* modeling

To identify potential regulators of sleep and cardiovascular physiology, and to reveal cross-tissue links between sleep disturbances and heart function, we queried large-scale GWAS for insomnia (*8*, *9*, *17*), and cardiovascular disease (*36–38*) for shared genomic signals and selected candidate genes for *Drosophila* modeling. We identified 13 genetic loci associated with insomnia that had previously been implicated in cardiovascular disease or its risk factors (Supp Table 1, 2, Fig. 1A). For example, at the 10q24.3 locus (Fig. 1B), insomnia and CAD (coronary artery disease) signals were in strong linkage disequilibrium (r^2^=0.85 between lead SNPs). At this locus, genetic associations (p<5×10^-8^) were observed with insomnia and difficulty awakening in the morning as well as with nine cardiovascular traits, including diastolic and systolic blood pressure, coronary artery disease, and myocardial infarction, reinforcing the genetic overlap between both diseases (Fig. 1C). Notably, at most genetic loci, genetic associations implicated the same proximal candidate genes for sleep and CVD traits, but lead signals were not necessarily in strong linkage disequilibrium, suggesting possible gene-level pleiotropy (Supp Fig 1). Thus the genetic signals may disrupt the same or nearby effector genes (Supp Table 3) to impact sleep and cardiovascular traits independently, or effector genes may influence sleep through cardiovascular dysfunction or vice versa. Based on proximity to association signals for insomnia and CVD traits, and gene expression in neuronal and cardiac tissues in both humans and flies, we selected 16 human genes at the 13 loci for *Drosophila* phenotypic screening (Supp Fig 2).

**Fig. 1.**
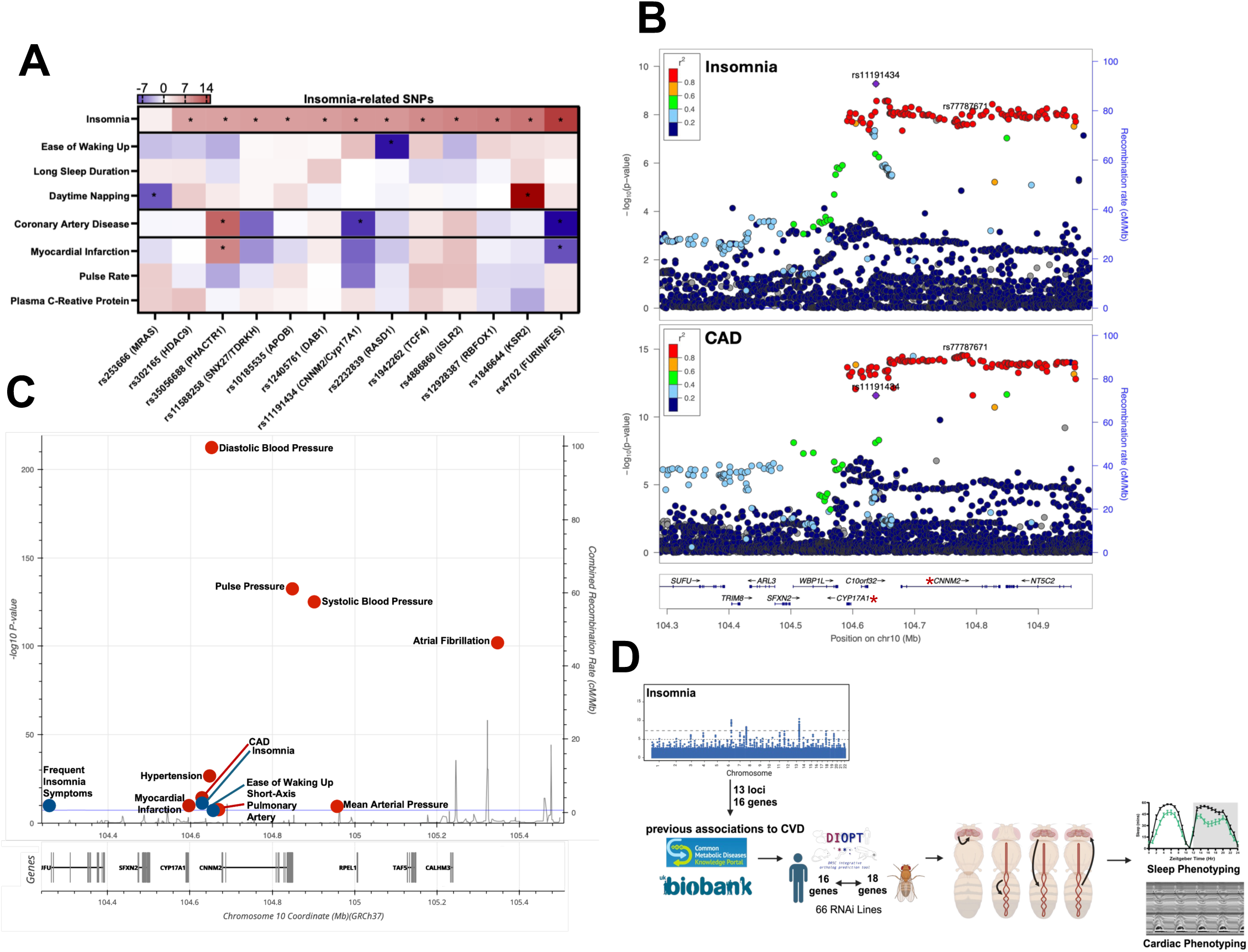
Shared genomic regions and GWAS signals between insomnia and CVD. (**A**) Associations of lead insomnia-related SNPs in various sleep and CVD-related GWAS studies obtained from the Knowledge Portal (37) and the literature (8). Color represents direction and value of Z-scores. Significant associations are labeled with an asterisk. (**B**) LocusZoom plots showing insomnia and CVD SNP association peaks with nearby genes. Purple diamond represents insomnia lead SNP (rs11191434) as LD reference. (**C**) GWAS associations meeting genome-wide significance (P<5×10-8, horizontal blue line) for sleep (blue dots) and cardiovascular (red dots) traits in the region near rs11191434 obtained from the Sleep and Cardiovascular Disease Knowledge Portal. Each point corresponds to a single trait in a single study. (**D**) Graphical Scheme showing study workflow.

From the 16 human candidate genes, we identified 18 corresponding *Drosophila* orthologs (Supp Table 4) and probed their roles in sleep and cardiac function by experimental downregulation of gene expression in a tissue-specific manner. We first knocked down the expression of the 18 genes (66 RNAi lines) using the *GAL4*/*UAS* system to express RNA interference (RNAi) constructs for each gene specifically in neurons using a pan-neuronal driver (*elav-Gal4)* and assessed their effects on sleep behavior and cardiac physiology. Then we expressed the RNAi constructs in the heart using *Hand-Gal4* and assessed their effects on cardiac physiology and sleep behavior. (Fig. 1D).

### Pan-neuronal knockdown of *Drosophila* orthologs impact behavioral sleep states

We tested if experimental knockdown of gene expression in neurons impacted sleep behavior using one week of continuous sleep and activity data recordings under normal light/dark conditions. We measured total sleep duration per 24-h interval, standard measures of sleep timing (including sleep duration during the day and night separately which can be regulated by distinct processes (*39*), standard measures of sleep quality (including sleep structure characterized by the number of sleep bouts (sleep episodes) and their average length), as well as sleep latency (time to fall asleep at night), wake after sleep onset (WASO; minutes active during the night after the first initiation of sleep), sleep discontinuity (1-minute-wakes; number of smallest possible wake periods), changes in morning or evening anticipation activity (*40*), and time spent in recently described *Drosophila* behavioral sleep states (*41*).

Robust alterations in sleep were observed for orthologs of 9/16 human genes; with evidence from 2 or more RNAi lines (Supp Table 5,6). Two primary phenotypes were observed among the set of genes tested. The first phenotype was marked by consistent decreases in total and/or nighttime sleep (Fig. 2A), along with some evidence of fragmented sleep (Supp Table 6). This “short sleeping” phenotype was observed for four fly genes, *Apoltp*, *Fur1*, *Cyp18a1* (Fig. 2D), and *da* (orthologs of *APOB*, *FUR*, *CYP17A1*, and *TCF4*, respectively). The second phenotype observed was marked by consistent increases in total and/or daytime sleep (Fig. 2A). This “long sleeping” phenotype was observed for three fly genes, *Ras85D*, *HDAC4*, and *Papi* (orthologs of *MRAS*, *HDAC9*, and *TDRKH*, respectively). Interestingly, a single RNAi line driving the knockdown of the scaffolding protein *Ksr*, which regulates RAS signaling, also resulted in increased sleep with *elav*>*Ksr^HMS00730^*flies experiencing 45% more minutes slept than controls (Fig. 2D). Together these data demonstrate that *Drosophila* modeling of genes associated with insomnia and CVD, recapitulate the short and long sleeping phenotypes that have been linked to increased risk of CVD (*42*).

**Fig. 2.**
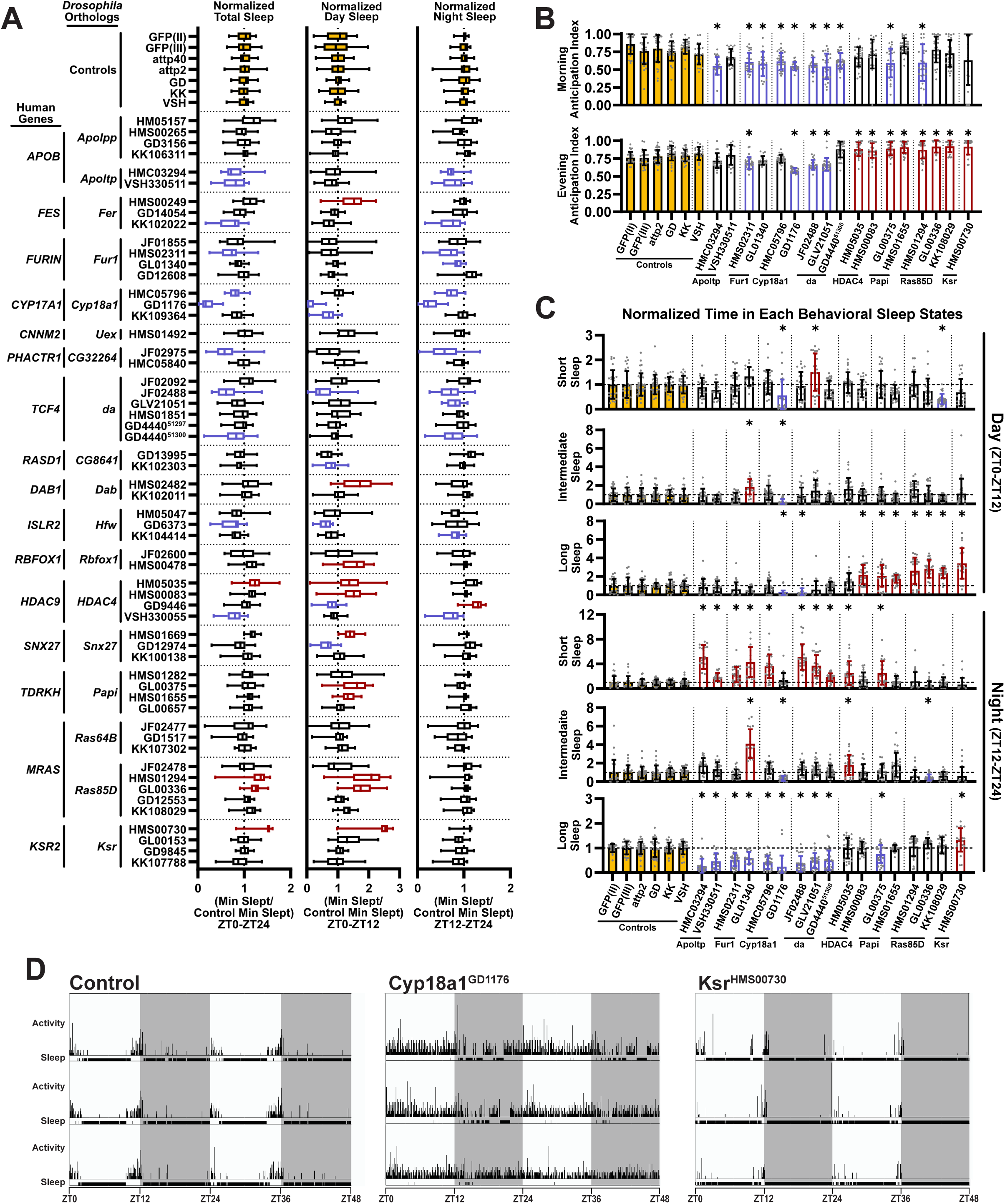
Neuronal-specific knockdown of insomnia- and CVD-associated genes impact behavioral sleep states and their timing. (**A**) Quantification of sleep quantity including total sleep (ZT0-ZT24), day (ZT0-ZT12) and night sleep (ZT12-ZT24). (**B**) Morning and evening anticipation index for a subset of knockdowns presenting with decreased or increased total, day, or night sleep. (**C**) Quantification of time spent in each behavioral sleep state during the day (top three graphs) and night (bottom three graphs) for the subset of knockdowns presenting with decreased or increased total, day, or night sleep. (**D**) Representative double-plotted actigraphy and sleep bouts for a control (elav-Gal4/+), elav-Gal4>Cyp18a1GD1176 (shortest sleeping line), and the longest sleeping line elav-Gal4>KsrHMS00730 (longest sleeping line). White areas indicate light periods and gray-shaded areas indicate dark periods of the 12:12 LD regime. Vertical black bars indicate summed activity counts for each 30-second period, Horizontal black bars indicate periods of sleep. Data was collected from at least two independent experiments. (**A**-**C**) Statistical significance was calculated by one-way ANOVA with Šidák’s multiple comparisons test to respective background matched controls of the underlying non-normalized data. Red indicates the sleep parameter for the indicated genotype is significantly increased. Blue indicates the sleep parameter for the indicated genotype is significantly decreased. Yellow indicates control groups. = significant at P < 0.05.

Among the short and long sleeping flies, distinct alterations in anticipatory behavior were also observed. Morning anticipatory behavior was consistently reduced upon knockdown of genes that resulted in the short sleeping phenotype, with some lines also resulting in decreased evening anticipation (Fig. 2B). Conversely, upon knockdown of genes that resulted in the long sleeping phenotype, all lines resulted in increased evening anticipation (Fig. 2B). While these changes are likely driven by increased sleep fragmentation in the short sleeping flies and increased sleep consolidation in the long sleeping flies, classic measures of sleep fragmentation and consolidation, concurrent alterations in bout number and bout length, did not reliably emerge among these two primary phenotypes (Supp Table 6). Therefore, we turned to assessing the impact of these genes on the timing of and time spent in the recently described *Drosophila* sleep states, which may better account for sleep fragmentation and consolidation compared to measures of average bout length (*41*). Analysis of the time spent in short sleep, intermediate sleep, and long sleep during the day and night revealed that all lines that produced the short sleeping phenotype resulted in significantly decreased time in the long sleep state during the night. Additionally, eight of the nine lines that produced the short sleeping phenotype significantly increased the time spent in the short sleep state at night (Fig. 2C), indicating overall more fragmented nighttime sleep compared to controls. Conversely, seven of the eight lines that produced the long sleeping phenotype resulted in significantly increased time in the long sleep state during the day with minimal changes to the time spent in other sleep states during the day (Fig. 2C), indicating an increase in consolidated daytime sleep.

Together, these characterizations highlight the diverse aspects of sleep regulated by *Drosophila* orthologues of insomnia-associated genes and reveal distinct sleep phenotypes in short sleeping and long sleeping marked by unique changes in sleep quantity, quality, and timing.

### Impact of neuronal-mediated short and long sleep phenotypes on *Drosophila* cardiac physiology

Given that both short and long sleep duration are associated with increased multivariable-adjusted risk of cardiovascular disease (*42*), we wanted to test whether pan-neuronal knockdown of *Drosophila* genes that result in the short sleeping or long sleeping phenotypes exerted cross-tissue effects on cardiac physiology. We selected two genes to represent the short sleeping phenotype, *Apoltp* and *da* (Fig. 2C), and two genes to represent the long sleeping phenotype, *Ras85D* and *Ksr* (Fig. 2C), and assessed their impacts on key cardiac physiological parameters of structure and function upon pan-neuronal knockdown.

Three-week-old male flies were dissected and imaged for assessment of key cardiac physiological parameters, including heart rate, arrhythmia index, diastolic and systolic intervals, diastolic and systolic diameters, and fractional shortening. Among the cardiac parameters measured, the lines that produced the short sleeping phenotype more consistently impacted cardiac physiology (Fig. 3A-F). While the long sleeping lines did not significantly alter heart rate, 3 of the 4 short sleeping lines resulted in significantly decreased heart rate compared to their respective controls (Fig. 3A). This bradycardic phenotype was primarily driven by an increase in the systolic interval (Fig. 3C). Similarly, while none of the long sleeping lines affected fractional shortening, three of the four short sleeping lines significantly lowered cardiac performance, as indicated by significantly lower fractional shortening (Fig. 3F). Although the long sleeping lines did not present with a consistent phenotype across all four lines, *elav>Ras85D^HMS01294^* resulted in a dilated phenotype indicated by increased diastolic and systolic diameters (Fig. 3C.,D). Additionally, none of the short sleeping or long sleep lines influenced arrhythmia prevalence (Supp Table 6). Together, these data suggest that disruption of insomnia-associated genes in the brain can influence cardiac performance, with those that drive fragmented and shorter sleep duration having a greater impact on cardiac parameters measurable in *Drosophila*.

**Fig. 3.**
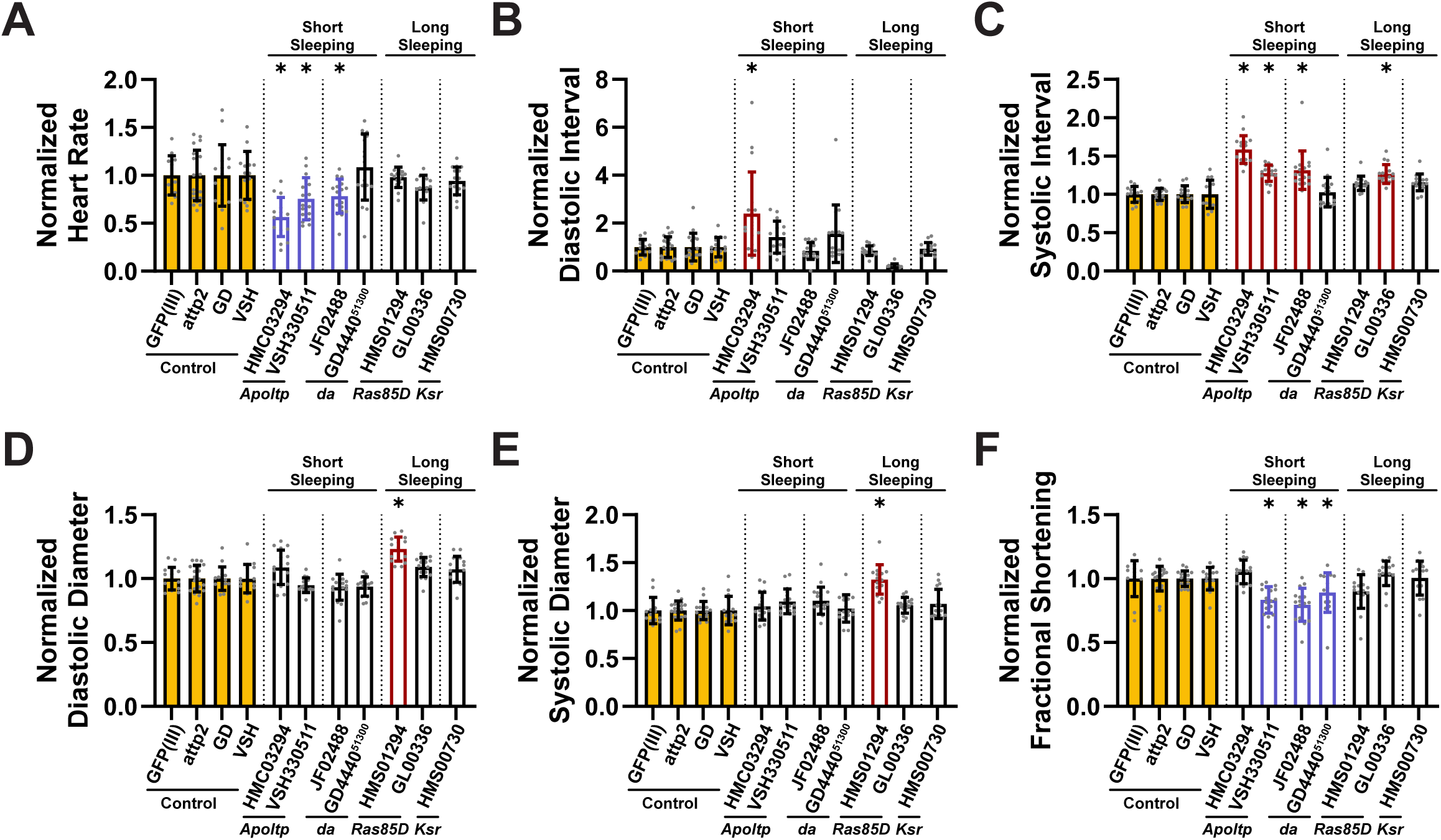
Disrupted neuronal expression of insomnia- and CVD-related genes resulting in short sleeping phenotypes, exhibit more impacts on heart function. Quantification of cardiac physiological parameters, including (**A**) heart rate, (**B**) diastolic and (**C**) systolic intervals, (**D**) diastolic and (**E**) systolic diameters, and (**F**) fractional shortening of three-week-old male flies with elav-Gal4-driven, neuronal-specific RNAi knockdown of representative short sleeping flies and representative long sleeping flies. (A-**F**) Parameters are normalized to each RNAi’s respective background matched control. Error bars represent the standard deviation. Red indicates lines with a statistically significant increase while blue indicates a line with a statistically significant decrease, = significant at P < 0.05. Data was collected from at least two independent experiments. Statistical significance was calculated by one-way ANOVA with Šidák’s multiple comparisons test to respective controls of the underlying non-normalized data.

### Cardiac-specific knockdown of *Drosophila* orthologs compromises cardiac physiology

While human candidate genes were also selected based on their previously reported associations with CVD traits and expression in human heart, it was important to assess if the genes played a role in regulation of cardiac structure and function in *Drosophila*, as opposed to other CVD-relevant tissues not well modeled in flies (*43–46*). Thus, we knocked down *Drosophila* orthologs of these genes specifically in the heart using *Hand-Gal4* and measured the previously mentioned cardiac parameters.

Consistent alterations in cardiac physiology were observed for five genes, supported by effects across at least two independent RNAi lines (Supp Table 7): *CG8641*, *CG32264*, *Uex*, *Ras85D*, and *da*, the *Drosophila* orthologs of *RASD1*, *PHACTR1*, *CNNM2*, *MRAS*, and *TCF4*. Two primary classes of phenotypes emerged among these genes. The first phenotype was characterized by altered cardiac rhythmicity and pacing. Knockdown of *CG8641* and *da* consistently decreased heart rate, while knockdown of *Uex* increased arrhythmia index (Fig. 4A,B), indicating disrupted cardiac rhythmicity. The second phenotype was characterized by altered cardiac contractility, with knockdown of *CG32264* and *Ras85D* increasing fractional shortening relative to controls (Fig. 4C). While broad shifts across all measured cardiac parameters were not observed, these reproducible changes across independent RNAi lines indicate that disruption of several CVD-associated genes produces specific and measurable defects in *Drosophila* cardiac physiology.

**Fig. 4.**
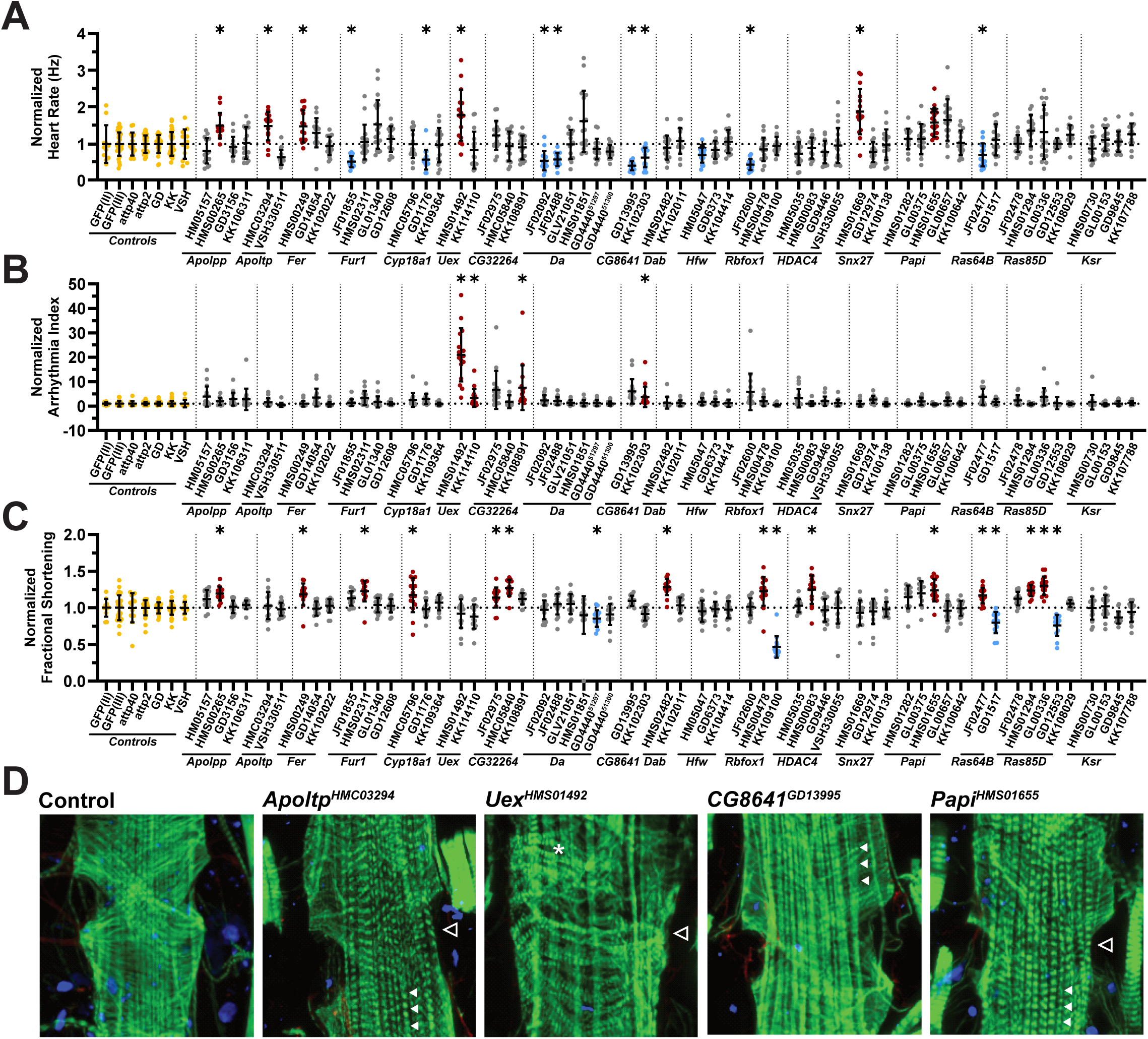
Cardiac-specific knockdown of insomnia- and CVD-associated gene impact measures cardiac rhythmicity and performance. Quantification of cardiac physiological parameters, including (**A**) heart rate, (**B**) arrhythmia index, (**C**) fractional shortening of three-week-old male flies with Hand-Gal4-driven, cardiac-specific RNAi knockdown. (**D**) Representative images showing actin-containing myofibrils (green), pericardin (red) and DAPI (blue). Open arrowhead indicates ostia defect, series of filled arrowhead indicates regions with increased visibility of longitudinal myofibrils, * indicates regions of disorganized circumferential myofibrils. (**A**-**D**) Parameters are normalized to the respective background matched control for each RNAi. Error bars represent the standard deviation. Red indicates lines with a statistically significant increase while blue indicates a line with a statistically significant decrease, yellow indicates control groups, = significant at P < 0.05. Data was collected from at least two independent experiments. Statistics significance was calculated by one-way ANOVA with Šidák’s multiple comparisons test to respective controls of the underlying non-normalized data.

To further assess whether these physiological abnormalities were accompanied by structural alterations in the heart, we characterized myofibrillar organization in representative lines showing distinct cardiac phenotypes. Control hearts displayed the expected circumferential organization of actin-containing contractile myofibrils. In contrast, cardiac-specific knockdown by *Uex^HMS01492^*, *Apoltp^HMC03294^*, *CG8641^GD13995^*, and *Papi^HMS01655^* resulted in evident disruption and loss of circumferential myofibrillar organization, which increased visibility of the non-contractile longitudinal myofibrils (Fig. 4D, filled arrowheads). Several of these lines also exhibited abnormal ostial morphology (Fig. 4D, empty arrowhead), suggesting broader defects in cardiac structural integrity. Although these structural analyses were performed on a subset of genes selected based on their cardiac phenotypes (Fig. 4D, Supp Fig. 3), the observed abnormalities support the conclusion that disruption of insomnia- and CVD-associated genes can impair both cardiac function and cardiac tissue organization.

Together, these characterizations of cardiac physiology and myofibrillar organization demonstrate that *Drosophila* orthologs of CVD-associated genes regulate multiple aspects of heart function and structure, revealing diverse cardiac phenotypes resulting from the disruption of genes implicated in both sleep and cardiovascular disease.

### Impact of altered cardiac performance on *Drosophila* behavioral sleep states

Next, to investigate whether cardiac dysfunction influences sleep behavior across tissues, we assessed sleep in the five *Drosophila* orthologs for which cardiac-specific knockdown produced robust and directionally consistent effects on at least one measured cardiac parameter in a minimum of two independent RNAi lines. Knockdown of *Uex* (*CNNM2*) by *Hand*>*Uex^HMS01492^* and *Hand*>*Uex^KK114110^*, both of which caused an increase in arrhythmia (Fig. 4B), significantly reduced morning anticipation (Supp Table 7). Knockdown of *Ras85D* (*MRAS*) by *Hand*>*Ras85D^HMS01294^*and *Hand*>*Ras85D^GL00336^*, which both increased fractional shortening (Fig. 4C), resulted in significantly fewer sleep bouts, increased morning anticipation, and a decrease in the time spent in the short sleep state driven by a decrease in daytime short sleep (Supp Table 7). Similarly, knockdown of *CG32264* (*PHACTR1*) with *Hand*>*CG32264^JF02975^* and *Hand*>*CG32264^HMC05840^*, both of which increased fractional shortening (Fig. 4C), resulted in decreased time in the short sleep state during the day. Interestingly, knockdown of *da* or *CG8641*, by *Hand*>*da^JF02092^*, *Hand*>*da^JF02488^*, *Hand*>*CG8641^GD13995^*, *Hand*>*CG8641^KK102303^*, all decreased heart rate (Fig. 4A), but only *da* resulted in increased nighttime sleep, characterized by more time spent in the long sleep state and a decrease in WASO (Supp Table 6). Altered sleep was not detected in either *CG8641* knockdowns in the heart. This difference between two genes with similar underlying cardiac phenotypes suggests a more complex relationship between the knockdown of these genes in the heart and their effects on sleep behavior.

Given the lack of consistently reproduced phenotypes among our set of genes, we set out to test if there was a clear influence of cardiac performance on sleep behavior in *Drosophila*. Since fractional shortening serves as a measure of cardiac contractility, we focused specifically on whether changes in fractional shortening were accompanied by reproducible alterations in sleep quantity or sleep structure across the tested lines. While consistent effects on total sleep duration were not observed across all lines, changes in fractional shortening were associated with distinct alterations in daytime and nighttime sleep. Among the 13 RNAi lines that increased fractional shortening, seven also exhibited increased nighttime sleep, whereas three of the four lines that decreased fractional shortening showed reduced daytime sleep relative to controls (Fig. 5A). To further assess whether cardiac dysfunction was associated with changes in sleep quality and fragmentation/consolidation, we analyzed time spent in the *Drosophila* sleep states. Among the lines that increased fractional shortening, six exhibited decreased time spent in the short sleep state together with increased time spent in the long sleep state, consistent with more consolidated sleep (Fig. 5B). However, some of these lines resulted in an increase in daytime long sleep, while others resulted in an increase in nighttime long sleep (Supp Table 7). In contrast, half of the lines that decreased fractional shortening showed the opposite pattern, with increased time spent in short sleep and decreased time spent in long sleep, indicative of more fragmented sleep (Fig. 5B). In these two cases, both demonstrated alterations in daytime sleep state but *Hand>Ras85D^GD12553^* also experienced alterations in nighttime sleep state (Supp Table 7). Together, these findings suggest that alterations in cardiac performance are associated with specific changes in sleep timing and quality in *Drosophila*.

**Fig. 5.**
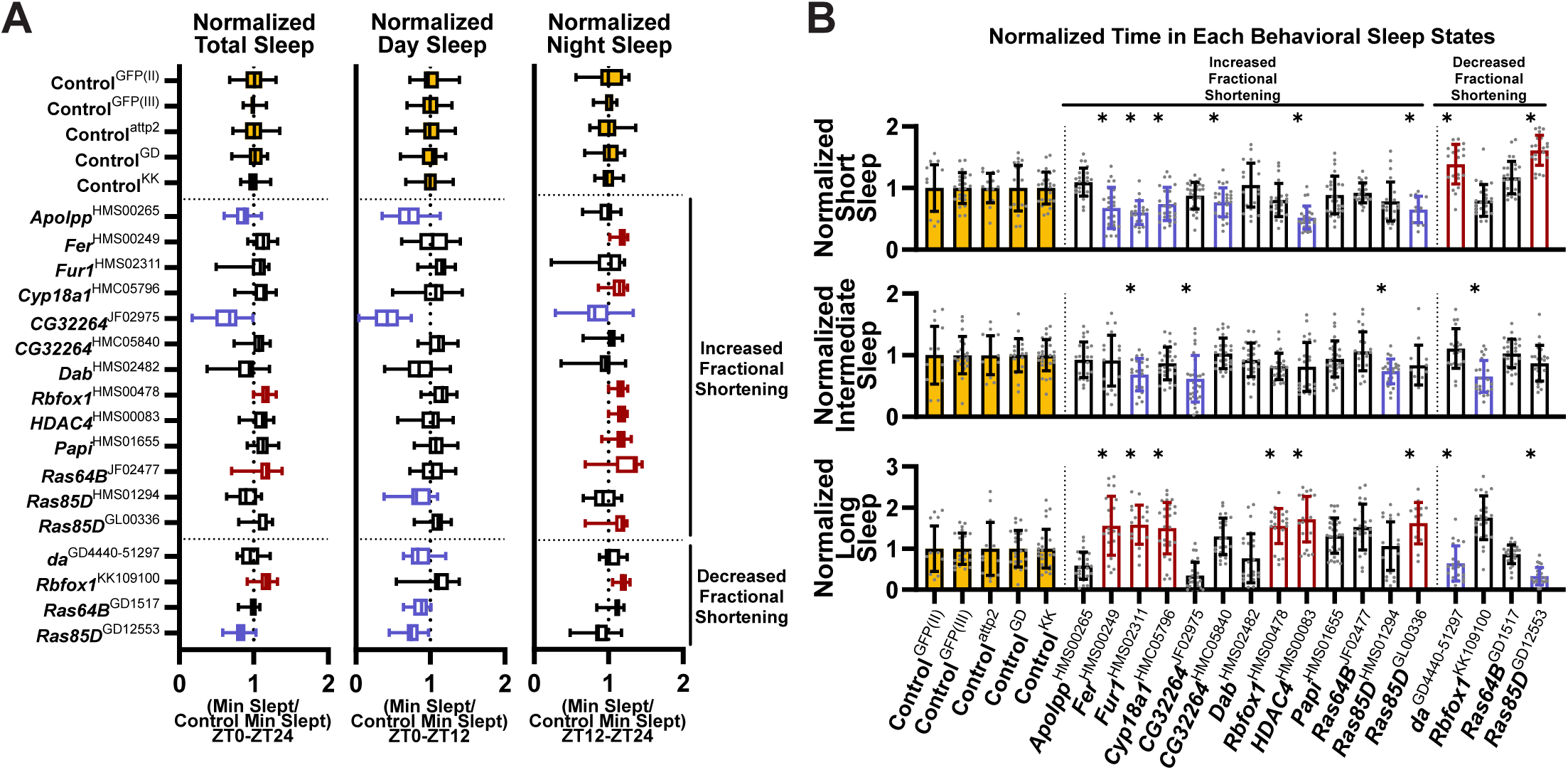
Disrupted cardiac expression of insomnia- and CVD-related genes that alter fractional shortening influence behavioral sleep states. (**A**) Quantification of sleep quantity including total sleep (ZT0-ZT24), day (ZT0-ZT12) and night sleep (ZT12-ZT24), and (**B**) average time spent in each behavioral sleep state over 24 hours for Hand-Gal4-driven, cardiac-specific knockdowns that resulted in significantly altered fractional shortening. Parameters are normalized to each RNAi’s respective background matched control. Error bars represent the standard deviation. Red indicates lines with a statistically significant increase while blue indicates a line with a statistically significant decrease, yellow indicates control groups, = significant at P < 0.05. Data was collected from at least two independent experiments. Statistical significance was calculated by one-way ANOVA with Šidák’s multiple comparisons test to respective controls of the underlying non-normalized data.

## DISCUSSION

*Drosophila* modeling of candidate gene orthologs from shared human insomnia and cardiovascular disease loci using neuronal and cardiac RNAi knockdown identified robust impact on sleep and cardiac phenotypes for several orthologs, demonstrating remarkable bi-directional tissue cross-talk. Specifically, neuronal knockdown of four fly orthologs resulted in short-sleeping flies with nighttime sleep fragmentation and three orthologs resulted in long-sleeping flies with increased consolidated daytime sleep. Short-sleeping fly lines also impacted cross-tissue cardiac physiology, primarily resulting in bradycardia and lower cardiac performance, consistent with links between short or poor-quality sleep and increased cardiovascular disease in humans. Conversely, disruption of five genes in the *Drosophila* heart led to defects in cardiac rhythmicity and pacing, cardiac contractility, or structural tissue organization, with wide-ranging effects on sleep. Notably, cardiac knockdown lines that altered cardiac performance also influenced long and short sleep states, with lines that increased fractional shortening, reflecting potential cardiac hypertrophy, increasing time spent in the consolidated long sleep state, whereas those that decreased fractional shortening, reflecting reduced cardiac performance, demonstrating reduced day sleep and increasing the time spent in the short sleep state.

Orthologs of nine of the 16 candidate human genes regulated sleep phenotypes in *Drosophila*, demonstrating both that *Drosophila* is a good model for investigating sleep disturbances seen in humans, and that our study implicates several effector genes at human insomnia genetic loci. In particular, genes with diverse molecular functions in a range of pathways led to short and fragmented sleep (orthologs of *CYP17A1*, *TCF4*, *APOB* and *FUR*). *Cyp18a1*, a *Drosophila* ortholog of *Cyp17a1*, encodes a protein that functions in ecdysone hormone signaling and the human ortholog encodes a critical enzyme for steroid biogenesis (*47*). *Da*, the *Drosophila* ortholog of *TCF4*, encodes a class 1 bHLH transcription factor that regulates the transcription of proteins involved in a variety of biological processes, including developmental processes and signaling response pathways (*48*, *49*). *Fur1*, the *Drosophila* ortholog of *FUR*, encodes an endoprotease that functions in cleaving proproteins within the secretory pathway. Targets of FURIN cleavage range from hormones and growth factors to cell-surface receptors (*50*). *Apoltp*, a *Drosophila* ortholog of *APOB*, encodes a lipid transport particle that facilitates the transfer of lipids from lipophorin to various tissues (*51*). Besides *da*/*TCF4*, which has previously been shown to regulate sleep in *Drosophila* (*27*) and mice (*52*), the remaining genes leading to short sleep have not been implicated in sleep regulation beyond genetic associations.

While their functions vary, *Cyp18a1*, *da*, *Fur1*, and *Apoltp* knockdowns all share a common phenotype marked by decreased sleep duration, namely nighttime sleep, and elevated sleep fragmentation indicated by decreased time in the long sleep state and increased time in the short sleep state during the night. While *Drosophila* behavioral sleep states have only recently been described (*41*, *53*), this short sleep duration and increased sleep fragmentation is similar to other *Drosophila* models of insomnia (*54–58*). While many short sleeping mutants in *Drosophila* have been uncovered through forward genetic screens, fewer examples exist starting from insomnia associations and revealing genetic drivers of short/fragmented sleep phenotypes (*27*, *59*). Our work expands the list of causal effector genes that can contribute to short sleep duration experienced by patients with insomnia by including *Cyp18a1*/*CYP17A1*, *Apoltp*/*APOB*, and *Fur1*/FUR.

Our modeling of insomnia-associated genes also revealed a sub-set of *Drosophila* orthologs that resulted in longer sleep during the day that is more consolidated (orthologs of *HDAC9*, *TDRKH*, and *MRAS*). *HDAC4*, the *Drosophila* ortholog of *HDAC9* with shared orthology to *HDAC4*/*5*/*7*, encodes a class IIa Histone deacetylase which functions in the regulation of gene expression (*60*). *Papi*, the *Drosophila* ortholog of *TDRKH*, encodes a protein that interacts with PIWI proteins and piRNA to regulate epigenetic modifications and silence transposons (*61*). *Ras85D*, a *Drosophila* ortholog of *MRAS*, encodes a small RAS GTPase involved in signaling pathways regulating tissue growth and development (*62*). Interestingly, an RNAi line targeting *Ksr*, which encodes a scaffold protein and positive regulator of the Ras/Raf-MAPK signaling cascade (*63*), resulted in the greatest increase in sleep and sleep consolidation from our modeling. While short sleep duration is strongly associated with insomnia, excessive daytime sleepiness is positively correlated with insomnia symptoms (*64*). Whether the long sleeping phenotype seen among the gene disruptions modeled in this work, represents excessive daytime sleepiness, increased napping, or hypersomnolence remains to be determined. However, increased and consolidated sleep has been seen previously in other work modeling disruption of insomnia-associated genes, including *PIG-Q/PIGQ* (*27*)*, ATPSynC/ATP5G1*, *Imp/IGF2BP1* (*29*), and modeling of loci associated with excessive daytime sleepiness, napping, ease of waking up, and chronotype, including *beat-1a*/*CADM2* (*65*) and *Alg10*/*ALG10*/*ALG10B* (*28*). Together *Drosophila* models of insomnia and insomnia-associated sleep traits indicate that long sleep is a common phenotype among some causal effector genes, which may highlight a possible subtype of insomnia with distinct underlying mechanisms.

Since both short and long sleep has been linked to increased risk of cardiovascular disease, such as myocardial infarction (*42*, *66*, *67*), we assessed metrics of cardiac physiology in *Drosophila* exhibiting the short or long sleep phenotypes caused by disruption of insomnia associated genes. We found that short sleeping flies more consistently led to impacts on cardiac function, namely decreased heart rate and reduced fractional shortening. While the long sleeping flies in our study experienced limited effects on cardiac function. However, previous *Drosophila* modeling of disrupted insomnia-associated genes, that resulted in longer sleep, have shown impacts on the heart. As mentioned previously, neuronal knockdown of *ATPSynC* and *Imp* both resulted in long sleeping flies. However, only *Imp* knockdown, which also resulted in sleep fragmentation, altered cardiac physiology (*29*). Similar to the impact of our short sleeping flies on the heart, the fragmented long sleep led to decreased heart rate and reduced fractional shortening (*29*). Importantly, in humans, sleep fragmentation index has been associated with carotid wall thickness (*68*), while in mice, acute sleep fragmentation drives the rapid expression of pro-inflammatory genes in the heart (*69*). Even though sleep duration differs between short and long sleeping flies, the presence of sleep fragmentation indicates that flies are spending less time in the long sleep state and more time in the short sleep state. One of the initial features in driving the cut off determination between the behaviorally defined *Drosophila* sleep states was the change in metabolic rate. Importantly, flies in the short sleep state have similar metabolic rates to those in the wake state. Furthermore, the long sleep state cut off is partly defined by a sustained stabilization of a lower metabolic rate (*41*, *70*). Therefore, flies experiencing sleep fragmentation and more time in the short sleep state maintain a heightened metabolic rate that could be placing excessive stress on the heart. Together these studies suggest that sleep fragmentation may be an important sleep metric in mediating the adverse effects of insomnia on the heart.

Cardiac-specific knockdown of *Drosophila* orthologs of insomnia- and CVD-associated genes revealed two broad classes of cardiac phenotypes: altered cardiac rhythmicity and altered contractility. The genes producing these phenotypes encode proteins with diverse molecular functions, yet several have previously established connections to cardiac biology that align with their observed phenotypes in our model. *CG8641*, the *Drosophila* ortholog of *RASD1* (also known as DEXRAS1), encodes a dexamethasone-inducible member of the Ras superfamily of small GTPases that modulates Gi/Go protein signaling (*71*). Notably, *RASD1* is highly expressed in the heart and its downregulation has been reported in atrial tissue under conditions of volume overload, consistent with a role in regulating cardiac G-protein signaling and with the reduced heart rate observed upon its cardiac knockdown (*72*). As previously mentioned, *da*, the *Drosophila* ortholog of TCF4, encodes a class I bHLH transcription factor with broad roles in development. Given the broad role of *da* in transcriptional regulation, its cardiac knockdown may influence heart physiology through altered expression of downstream genes required for cardiomyocyte function (*48*). *Uex*, the *Drosophila* ortholog of *CNNM2*, encodes a magnesium transporter involved in cellular Mg^2+^ homeostasis (*73*). Loss of CNNM2 function leads to hypomagnesemia, which is known to increase arrhythmia risk through disruption of Mg^2+^-dependent regulation of calcium handling in cardiomyocytes, consistent with the increased arrhythmia index observed upon *Uex* cardiac knockdown (*73–76*). *CG32264* and *Ras85D*, orthologs of *PHACTR1* and *MRAS* respectively, both increased fractional shortening when knocked down in the heart. *PHACTR1* is one of the most commonly identified loci in GWAS studies of coronary artery disease and myocardial infarction and has been implicated in regulation of actin cytoskeletal organization in cardiomyocytes, suggesting that disruption of *CG32264* may alter contractile architecture and cardiac function (*77*, *78*). For *MRAS*, gain-of-function mutations have been identified in Noonan syndrome with cardiac hypertrophy, acting through enhanced *MRAS* signaling and downstream Ras/MAPK pathway activity (*79*, *80*). The increased fractional shortening observed upon *Ras85D* cardiac knockdown likely reflects altered Ras/MAPK signaling dynamics and changes in contractile regulation, consistent with the broader role of this pathway in cardiac growth and remodeling (*81*). Together, the diverse molecular functions of these genes and their alignment with known cardiac biology underscore the utility of *Drosophila* cardiac-specific knockdown as a platform for functional validation of CVD-associated candidate genes and for dissecting their roles in heart physiology.

Disruption of cardiac function through heart-specific knockdown also had wide-ranging effects on sleep behavior, revealing a cross-tissue influence of cardiac performance on sleep regulation. Notably, alterations in fractional shortening were associated with distinct changes in sleep timing and architecture. This relationship between cardiac performance and sleep architecture is consistent with previous findings, where cardiac dysfunction has been shown to alter sleep structure, and extends these observations by suggesting that heart-to-brain communication pathways linking cardiovascular state to sleep regulation may be evolutionarily conserved (*22*, *82*, *83*). The presence of sleep fragmentation across lines with reduced cardiac performance is particularly noteworthy as short-sleeping neuronal knockdown lines also exhibited reduced fractional shortening. This brain-to-heart and heart-to-brain communication suggests a possible feedback loop between the brain and heart centered around short sleep and decreased cardiac performance. Recent work in mammalian models further supports bidirectional heart-brain communication, demonstrating that cardiac injury can alter sleep through immune-mediated signaling pathways and that sleep disruption can worsen cardiac inflammation and recovery (*22*). Two non-mutually exclusive mechanisms may underlie the link between cardiac dysfunction and sleep fragmentation. First, as previously mentioned, flies in the short sleep state maintain a metabolic rate comparable to wakefulness, whereas the long sleep state is characterized by a sustained reduction in metabolic activity. Cardiac dysfunction may therefore impose a metabolic burden that biases flies away from sustained long sleep and toward shorter sleep states. Second, inflammatory signaling may contribute to heart-to-brain communication that shapes sleep behavior (*84*). NF-κB signaling has been shown to be activated in cardiomyocytes during cardiac stress and dysfunction (*85*, *86*). Notably, NF-κB also regulates innate immunity and sleep, with immune activation increasing sleep through NF-κB-dependent pathways, suggesting a potential mechanistic link between cardiac inflammation and sleep behavior. (*19*, *87–90*). Together, these findings raise the possibility that cardiac dysfunction in our knockdown lines could promote inflammatory signaling that in turn disrupts sleep architecture, a hypothesis that warrants direct experimental testing. Conversely, the increased sleep consolidation observed in lines with elevated fractional shortening may reflect altered compensatory sleep regulation in response to changes in cardiac workload, consistent with reported observations where elevated cardiac demand is associated with increased sleepiness (*22*). Together, these findings suggest that alterations in cardiac performance bidirectionally influence sleep architecture in *Drosophila*, and that both metabolic and inflammatory pathways represent plausible mechanisms linking the cardiac state to sleep-regulatory circuits.

Major strengths of our study lie in leveraging the genetic versatility of *Drosophila* to functionally validate candidate genes from human GWAS loci for both insomnia and cardiovascular disease in a single model system, enabling tissue-specific dissection of gene function in neurons and the heart. The use of both neuronal and cardiac RNAi knockdown allowed for the direct examination of bidirectional heart-brain cross-talk, revealing conserved pathways between cardiac performance and sleep regulation. Furthermore, this study incorporates the newly defined *Drosophila* behavioral sleep states (*41*), allowing for a more nuanced characterization of sleep architecture beyond simple duration metrics, including sleep consolidation and fragmentation. In addition, the range of sleep phenotypes observed across control lines provides a robust baseline against which gene-specific effects could be reliably detected. Finally, the convergence of findings across multiple genes and pathways, including Ras signaling, highlights the utility of *Drosophila* as a platform for uncovering conserved mechanisms underlying the comorbidity of insomnia and cardiovascular disease.

Despite these strengths, there are several limitations to our study that must be acknowledged when interpreting the results. The insomnia phenotype in the UKB used for prioritization of genes was based on self-reported symptoms of trouble falling asleep and maintaining sleep, which is less severe than an insomnia diagnosis that also relies on three month chronicity of symptoms and significant daytime impairment. Orthology determination tools have expanded since the inception of this study. In some cases, updated prediction tools have revealed additional higher ranking *Drosophila* orthologs for some human genes, with closer orthology than the genes selected in this study. The *Drosophila* modeling in this study focused on the impact of loss of function through tissue specific knockdown. However, for some genes insomnia- and CVD-associated variants may result in increased expression or gain-of-function, which were not assessed in this study. Additionally, phenotyping of sleep upon disruption of insomnia-associated genes was carried out under neuronal disruption conditions. While neurons compose a majority of the tissue of the brain, glial cells have been shown to contribute to the regulation of sleep. It is possible that some of the symptoms experienced by insomnia and CVD patients are driven by disruption of gene functions in glia. Furthermore, for the phenotyping of CVD-related traits it is important to acknowledge the limitations of the *Drosophila* model. Due to the fly open circulatory system, the effects of CVD-associated genes on vascular tissue cannot be assessed. For some CVDs, such as CAD, vascular effects are likely important for impacts on both sleep and the heart.

In conclusion, our study offers novel genetic evidence linking insomnia and cardiovascular disease, providing valuable insights into potential therapeutic targets for mitigating both conditions. We establish a list of insomnia- and CVD-associated genes and utilize tissue-specific genetic disruptions of their *Drosophila* orthologs to understand their cell-autonomous and non-cell-autonomous roles in regulating sleep and cardiac physiology. Through these characterizations, we find heterogeneity in the effects of the insomnia- and CVD-associated orthologs on sleep and cardiac physiology. Nonetheless, we uncover evidence indicating that non-cell-autonomous impacts of cardiac disruptions on sleep and neuronal disruptions on cardiac function are linked by excessive sleep. While cardiac knockdowns leading to broad yet diverse changes in cardiac physiology increase nighttime sleep, neuronal knockdowns that result in increased sleep tend to result in tachycardic heart phenotypes. Genetic correlation analysis of human GWAS data found that nearly all traits tested related to long sleep significantly correlated with heart failure, coronary artery disease, and myocardial infarction. Further investigation is needed to refine our understanding of the relationship between insomnia and CVD, but this work suggests that pathways implicated in increased and excessive sleep will provide valuable insight.

## MATERIALS AND METHODS

### LocusZoom plots and multi-trait associations for insomnia- and CVD-related SNPs

LocusZoom plots for insomnia- and CVD-related SNPs were generated using LocusZoom v1.3. Nearby associations for one representative SNP, rs11191434, were obtained from the Sleep and Cardiovascular Disease Knowledge Portal (Sleep Disorder Knowledge Portal)(*37*). Results were plotted using LDassoc in LDLink (*91*). Z scores for top insomnia SNPs from sleep and CVD-related GWAS were also obtained from the Sleep and Cardiovascular Disease Knowledge Portal after alleles were aligned.

### Drosophila husbandry

All *Drosophila* were raised and maintained on standard cornmeal fly food media with a 12h/12h light/dark cycle at 25°C with humidity between 50-70%. Strains used in this study were obtained from Bloomington *Drosophila* Stock Center (BDSC) or Vienna *Drosophila* Resource Center (VDRC), unless otherwise stated, and include: *elav-Gal4* (BDSC#: 458), *Hand-Gal4* (Gift from Eric Olson)(*92*), control lines: *Oregon-R-P2* (Oregon-R; BDSC#: 2376), *w^1118^* (BDSC#: 5905), *UAS-GFP RNAi* (II) (BDSC#: 41552) , *UAS-GFP RNAi* (III) (BDSC#: 41553) , *attp40* (BDSC#: 36304), *attp2* (BDSC#: 36303), *GD* (VDRC#: 60000), *KK* (VDRC#: 60100), *VSH* (VDRC#: 60200), Candidate gene RNAi stock numbers are listed in Supp Table 5 with complete genotypes for all lines used. RNAi stocks of the VRDC KK collection inserted into the 40D site were excluded from *elav-Gal4*-driven knockdown to avoid complications stemming from ectopic *tiptop* expression impacting wing development and therefore behaviors related to grooming or impaired locomotion (*93*).

### *Drosophila* sleep-wake behavior monitoring and analysis

*Drosophila* sleep-wake behavior was monitored through quantification of infrared beam breaks using the Drosophila Activity Monitoring (DAM) system (Trikinetics, Waltham MA). Individual 3-6-day-old male flies (cell-autonomous characterization; *elav-Gal4*-driven) or 17-20-day-old male flies (non-cell-autonomous; *Hand-Gal4*-driven) were loaded into glass tubes containing standard cornmeal fly food media. Flies were allowed to acclimate to monitoring conditions (12h/12h light/dark cycle at 25°C with humidity between 50-70%) for two days, then activity data was collected for five days. Flies that did not survive the full five days of data collection were not analyzed. All measurements represent the average of a given metric over the five days of monitoring. Activity data was analyzed using ClockLab and RStudio. Sleep in *Drosophila* is defined as any period of behavioral quiescence five minutes or longer as previously described (*23*, *24*). Total sleep included sleep during the full 24-hour period (ZT0-ZT24), day sleep included sleep between ZT0 and ZT12, and night sleep included sleep between ZT12 and ZT24 (*23*, *24*). Anticipatory behavior was calculated as previously described (*94*). Briefly, morning anticipation was calculated as the sum activity counts three hours prior to the light period divided by the sum activity counts six hours prior to the light period, evening anticipation was calculated as the sum of activity counts three hours prior to the dark period divided by the sum activity counts six hours prior to the dark period. Sleep latency was calculated as the minutes from lights off (ZT12) until the first bout of sleep. If flies were already asleep at lights off, no sleep latency measure was recorded. Bout number includes the total number of bouts during the full 24-hour period (ZT0-ZT24), and bout length represents the average length in minutes of all sleep bouts during the 24-hour period (ZT0-ZT24). Sleep discontinuity was calculated as previously described (*28*); sleep discontinuity was measured as the number of one-minute wakes during the 24-hour period. Time in each behavioral sleep state was calculated as previously defined (*41*). Sleep bouts were stratified into those lasting 5-29 minutes (short sleep), 30-59 minutes (intermediate sleep), and 60 or more minutes (long sleep). All minutes within a bout were attributed to the sleep state that the sleep bout was classified as. Cumulative time in the short sleep, intermediate sleep, or long sleep state were calculated for the full 24 hours, day, and night as defined for total sleep. At least 20 flies (cell-autonomous characterization; *elav-Gal4*-driven) or 15 flies (non-cell-autonomous; *Hand-Gal4*-driven) of each genotype were tested. Males were utilized to eliminate complications of egg laying and larval wandering causing erroneous beam breaks during the last days of data collection. DAM system data was collected and analyzed in RStudio using a custom code. RStudio scripts and methodology can be found on https://github.com/jameswalkerlab/Gill_et.al. One-way ANOVA with Šídák’s multiple comparisons test for DAM system data were performed using GraphPad Prism.

### Cardiac physiological analyses of semi-intact *Drosophila* hearts

Three-week-old male progeny of *Hand-Gal4* (cell-autonomous) or *elav-Gal4* (non-cell-autonomous) with control and RNAi lines of different genes were collected, and semi-intact hearts were prepared (*95*, *96*). Direct immersion optics were used in conjunction with a digital high-speed camera to record 30-second videos of beating hearts; images were captured using HC Image (Hamamatsu Corp.). Cardiac function for at least 30 flies was analyzed semi-automatic optical heartbeat analysis (SOHA) software to quantify heart rate, heart period, diastolic and systolic diameters, diastolic and systolic intervals, cardiac rhythmicity, and fractional shortening (*95*, *96*).

### Cytological studies of adult hearts

Dissected hearts from three-week old adults were treated with 5 mM EGTA in hemolymph then fixed with 4% paraformaldehyde in PBS as previously described(*95*). Fixed hearts were stained with anti-Pericardin antibody overnight (5µg/ml, 1:10; Developmental Biology Hybridoma Bank, University of Iowa) followed by Alexa488-phalloidin for 30 min (1:1000, U0281, Abnova). Diamond Antifade Mountant with DAPI was used to mount samples. Confocal images were acquired using a Nikon A1R HD microscope at 10X for pericardin quantification and 20X for representative images for phalloidin staining. Quantification of the pericardin area was done as previously described^65^.

### Statistical analysis

For all quantitation, statistical significance was determined using one-way analysis of variance (ANOVA) followed by Šidák’s post hoc test to determine significance between groups for sleep and cardiac physiological parameters compared to their respective controls. For pericardin quantification, statistical significance was determined using one-way ANOVA followed by Dunnett’s post hoc test to compare groups to the control. Bar graphs show mean ± SD. All statistical analyses were performed with GraphPad Prism 9. P-values indicated in figures are from one-way ANOVA followed by Šidák’s post hoc test of raw (non-normalized) data.

## Acknowledgments

We would like to thank Dr. Olson for supplying the *Hand-Gal4* driver stock. Stocks obtained from the Bloomington *Drosophila* Stock Center, and Vienna *Drosophila* Resource Center were used in this study. We would like to thank Stephanie Bouley for her help in preparing the manuscript and proof-reading. Figure 1D was created with BioRender.com.

## Funding

National Institutes of Health grant T32HL007901 (T.R.M.)

American Heart Association grant 23PRE1020631 (F.A.)

National Institutes of Health grant R01HL146751 (G.C.M., J.A.W., R.S.)

## Author contributions

Conceptualization: T.R.M., F.A., G.C.M., J.A.W., R.S.

Methodology: T.R.M., F.A., G.C.M., J.A.W., R.S.

Investigation: T.R.M, F.A., D.P., S.M., L.O., M. M.

Visualization: T.R.M, F.A.

Supervision: G.C.M., J.A.W., R.S.

Writing—original draft: T.R.M, F.A.

Writing—review & editing: G.C.M, J.A.W. and R.S.

## Competing interests

The authors declare they have no competing interests.

## Notes

### Competing Interest Statement

R.S. is a founder and stockholder of Magnet Biomedicine. The other authors declare no competing interests.

### Summary of Updates

The manuscript was fully overhauled based on findings from a more in-depth analysis of Drosophila sleep.

