## Supplemental Figures 1-3 and Supp Table Legends for "Bidirectional links between sleep regulation and cardiac physiology revealed by *Drosophila* modeling of genes at insomnia and cardiovascular disease loci"

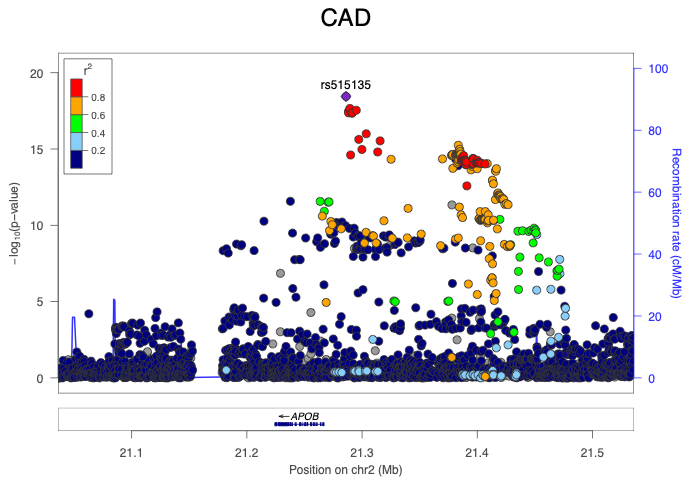

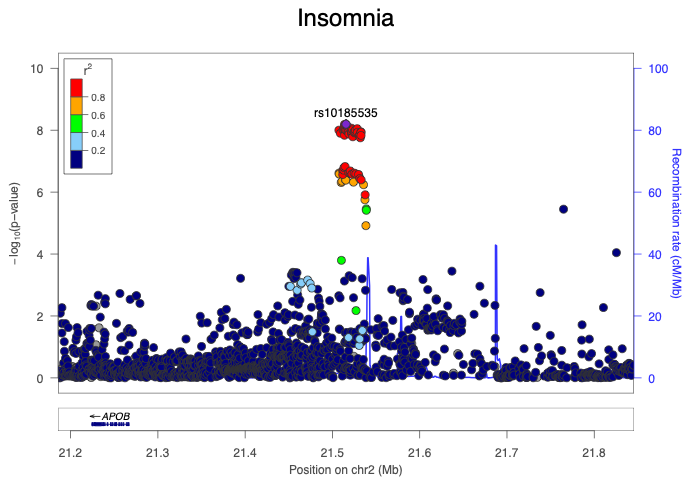

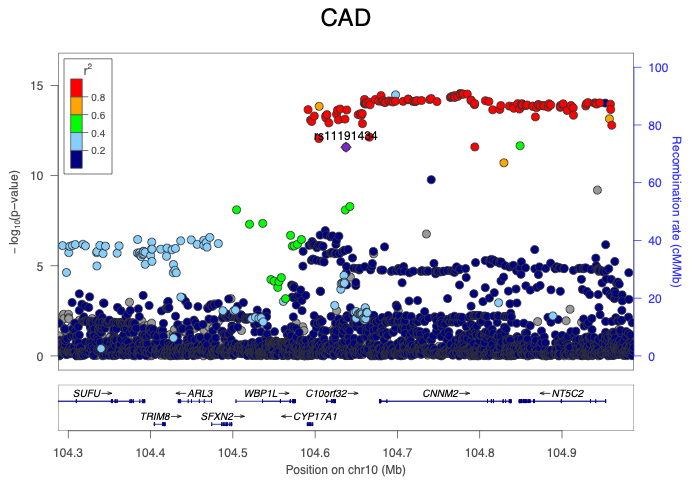

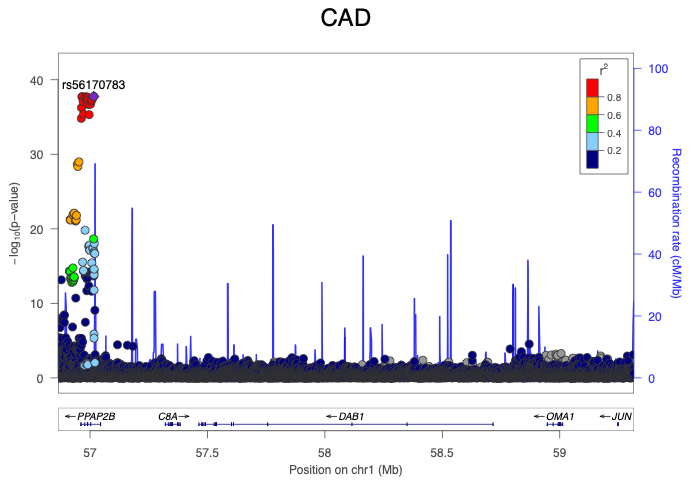

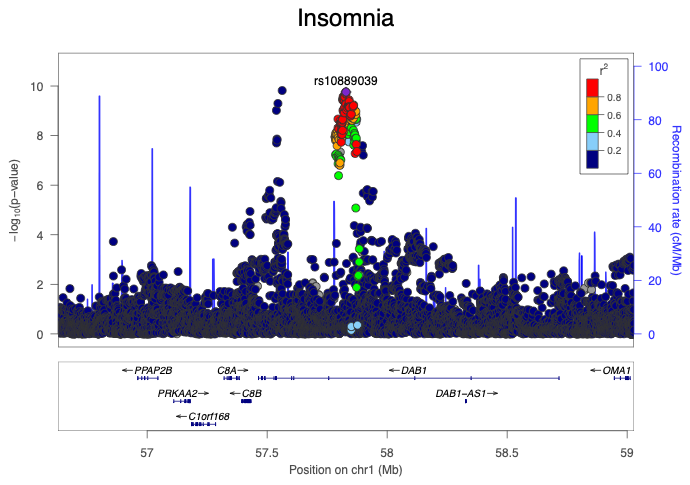

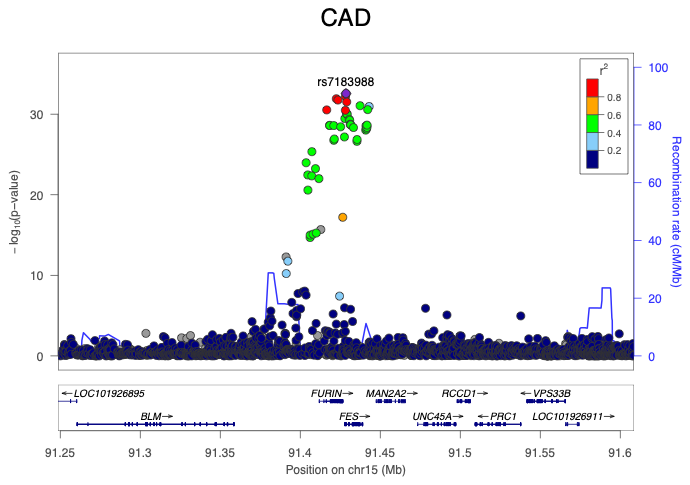

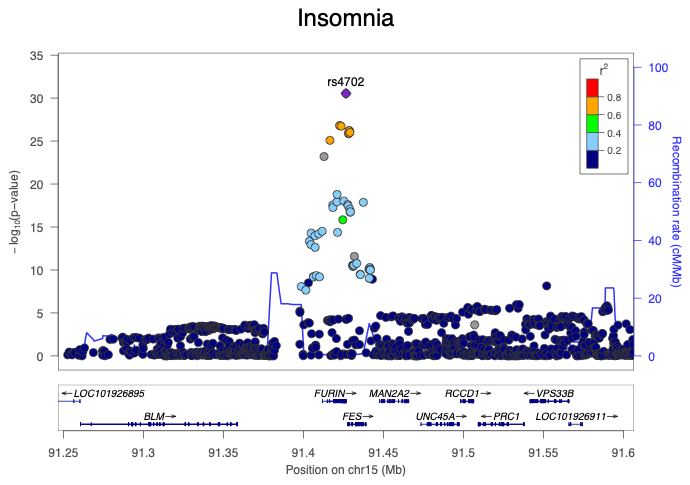

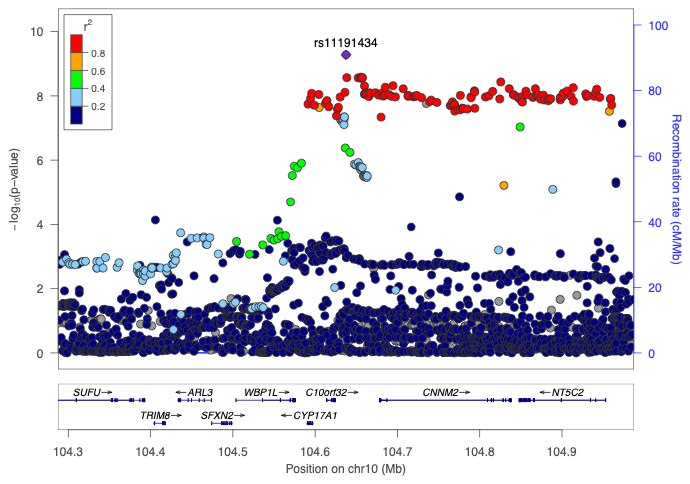

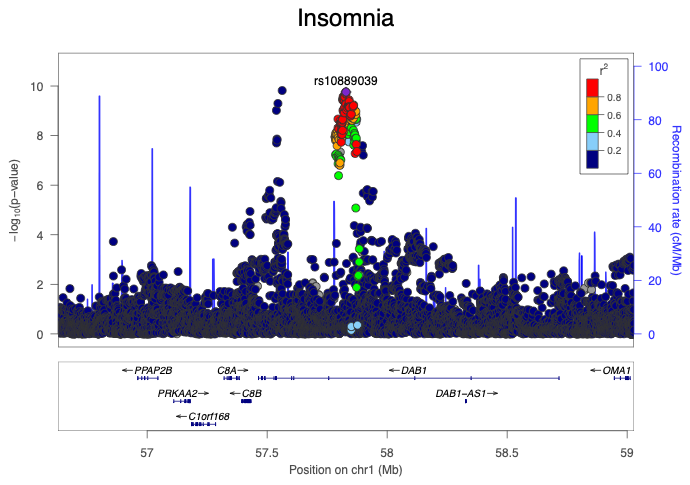

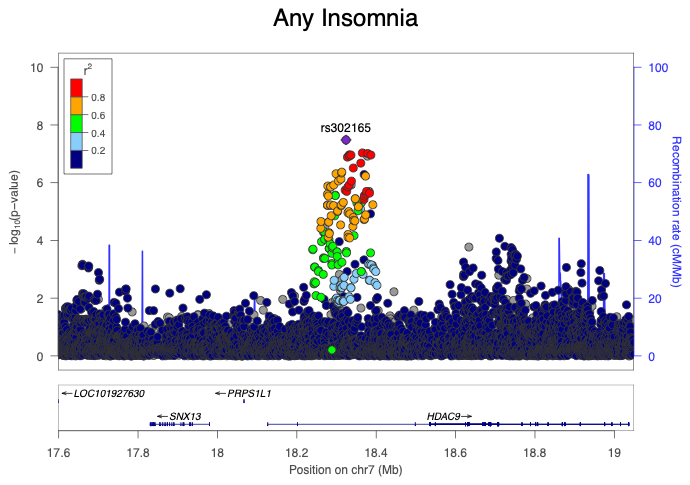

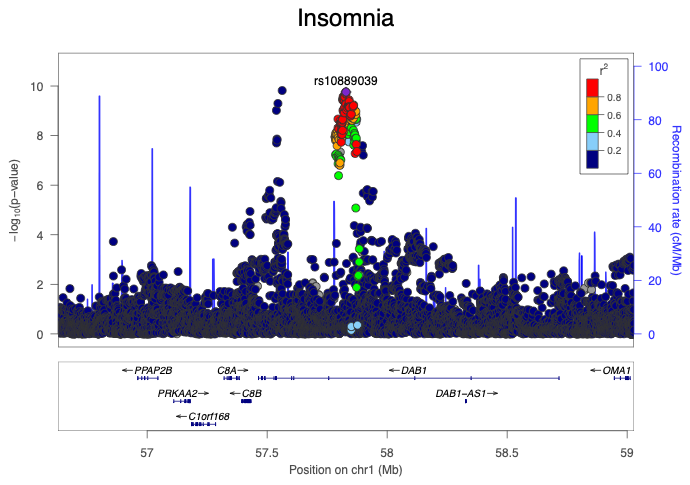

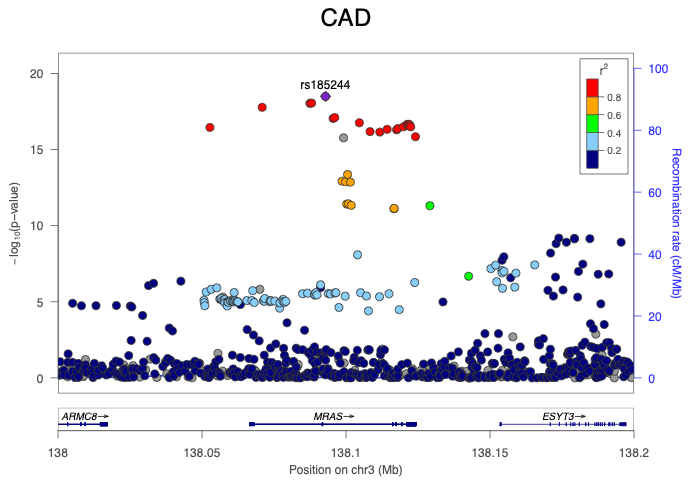

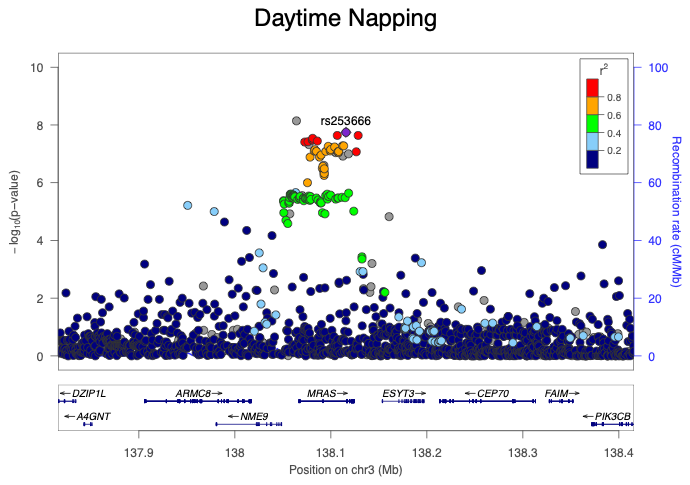

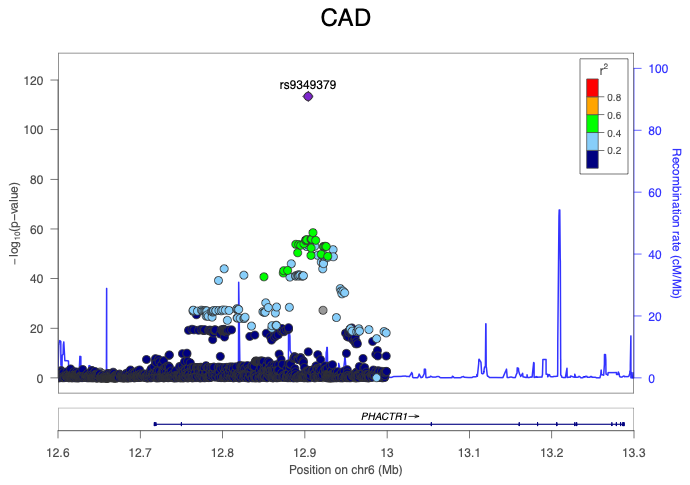

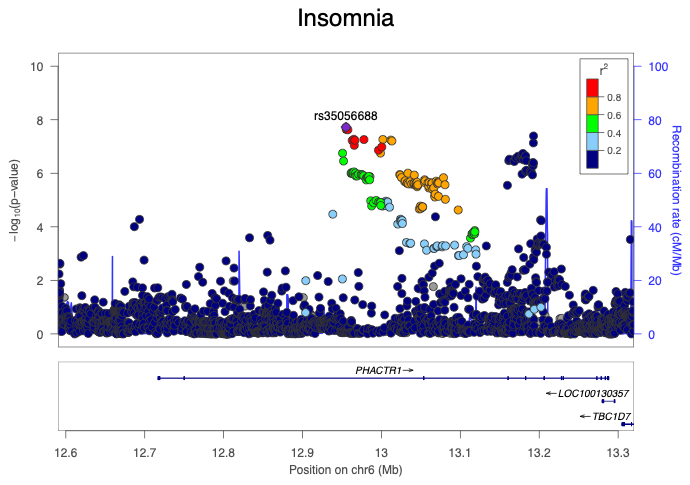

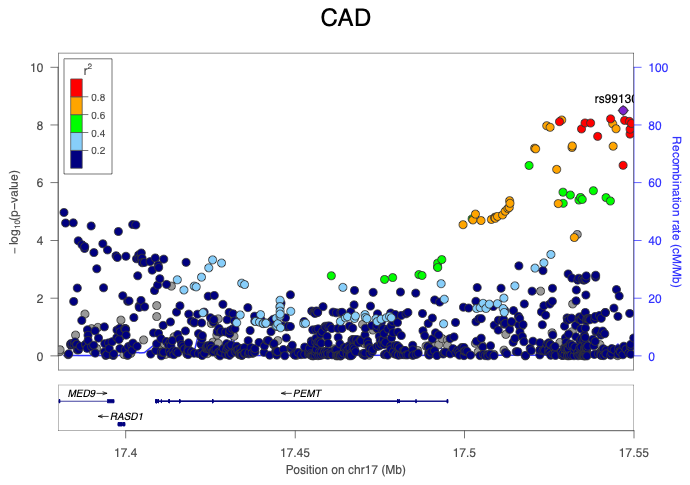

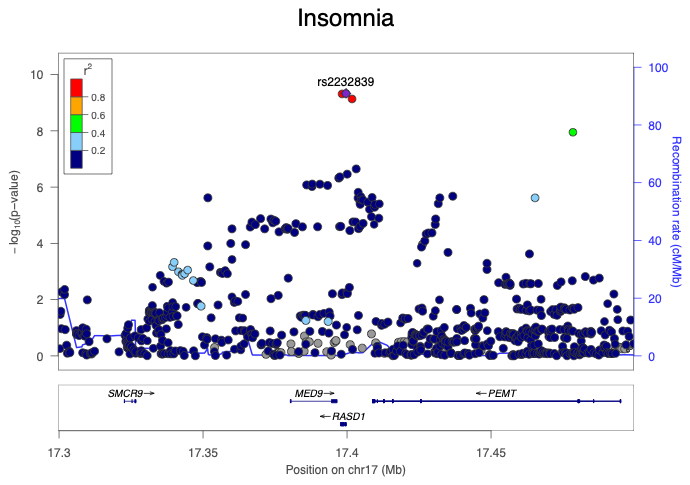

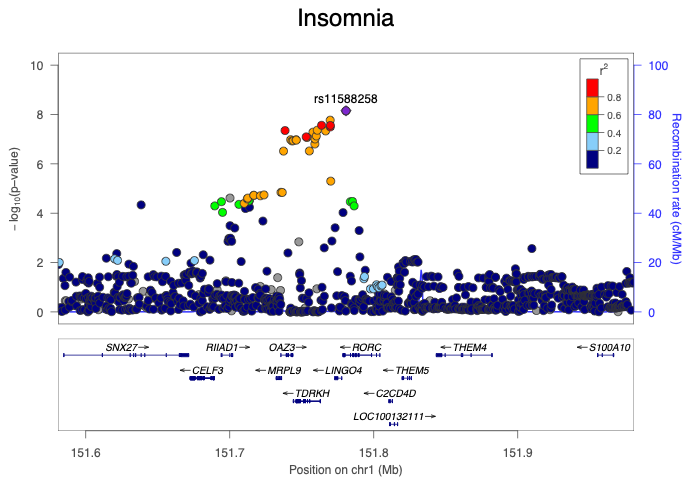

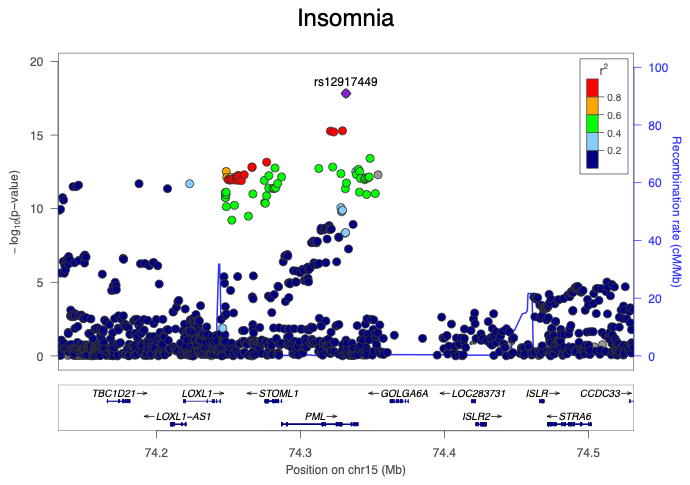

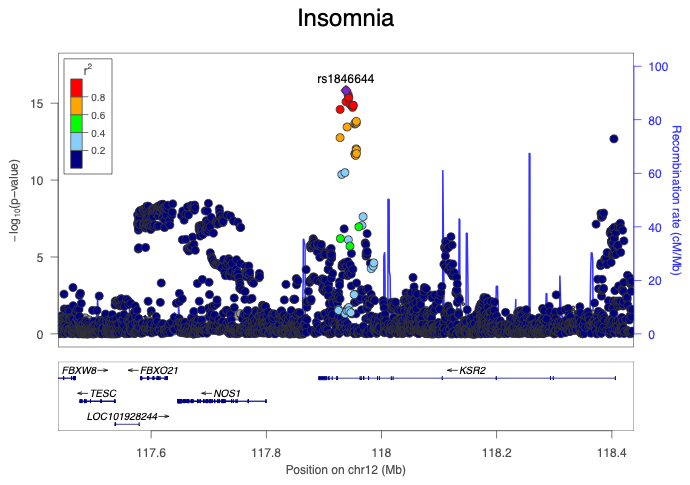

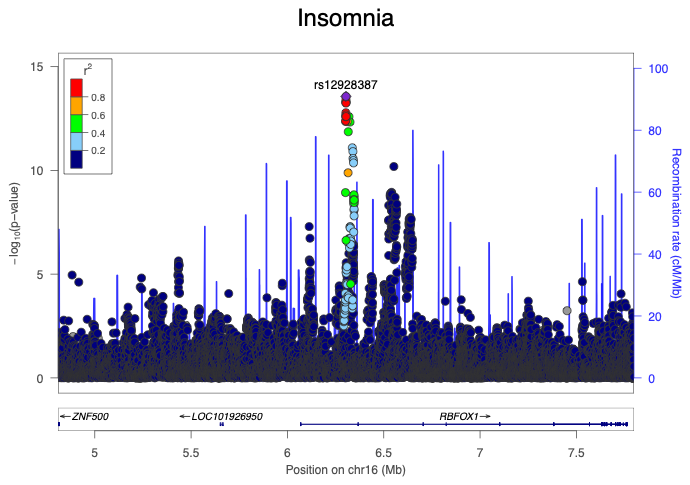

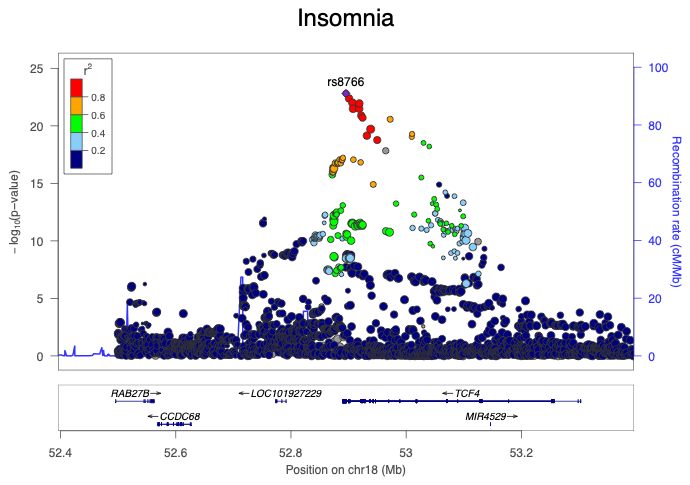

Fig. S1. LocusZoom regional association plots of the genome-wide significant loci associated with cardiovascular disease and sleep traits.

The lead single-nucleotide polymorphism (SNP) is indicated by a purple diamond, and surrounding variants are colored according to their linkage disequilibrium (LD; r²) with the lead SNP. The lower panel displays nearby genes within each locus.

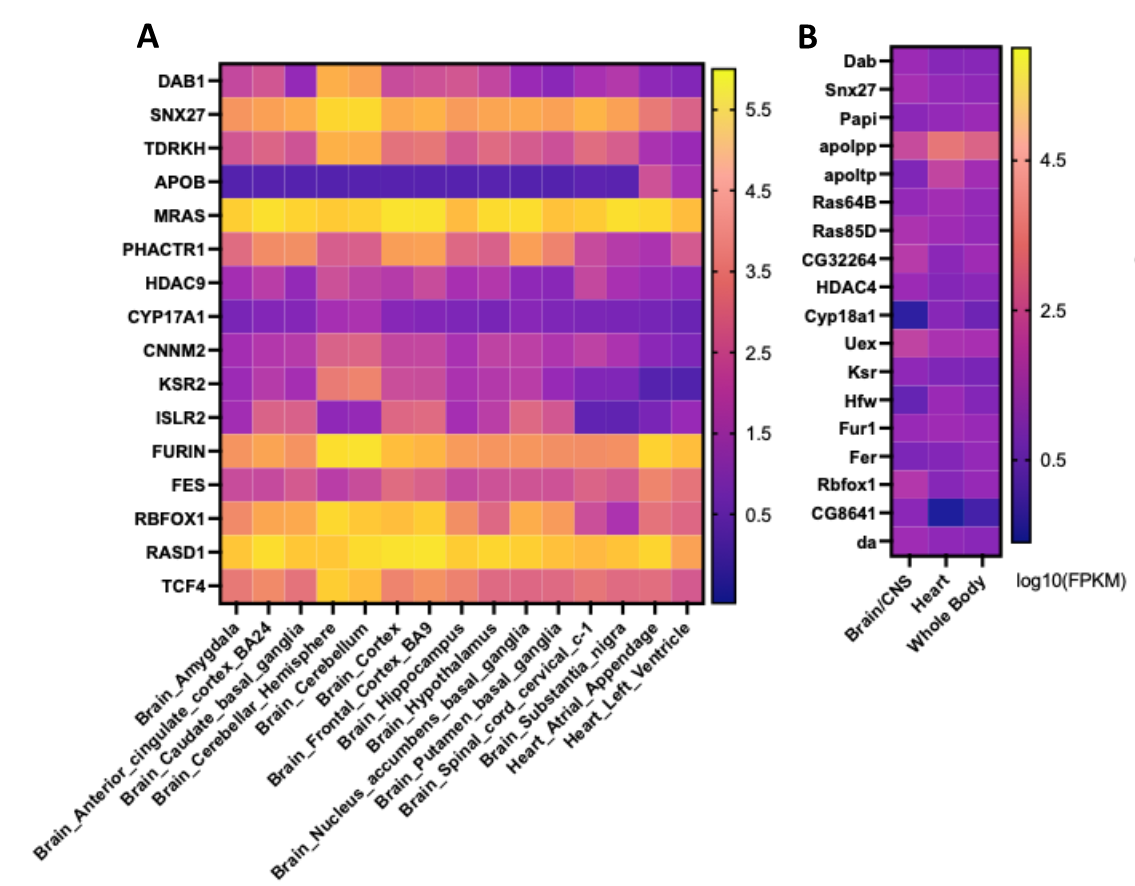

Fig. S2. Expression profiles in the brain and heart tissue of priotized insomnia- and CVD- related genes.

(**A**) Human gene expression across selected brain regions and heart tissues obtained from GTEx. (**B**) Expression of *Drosophila* orthologs across whole body, central nervous system (CNS)/brain, and heart tissues obtained from FlyAtlas. Color intensity represents log10-transformed expression values, shown as transcripts per million (TPM) for human tissues and fragments per million (FPM) for *Drosophila* tissues.

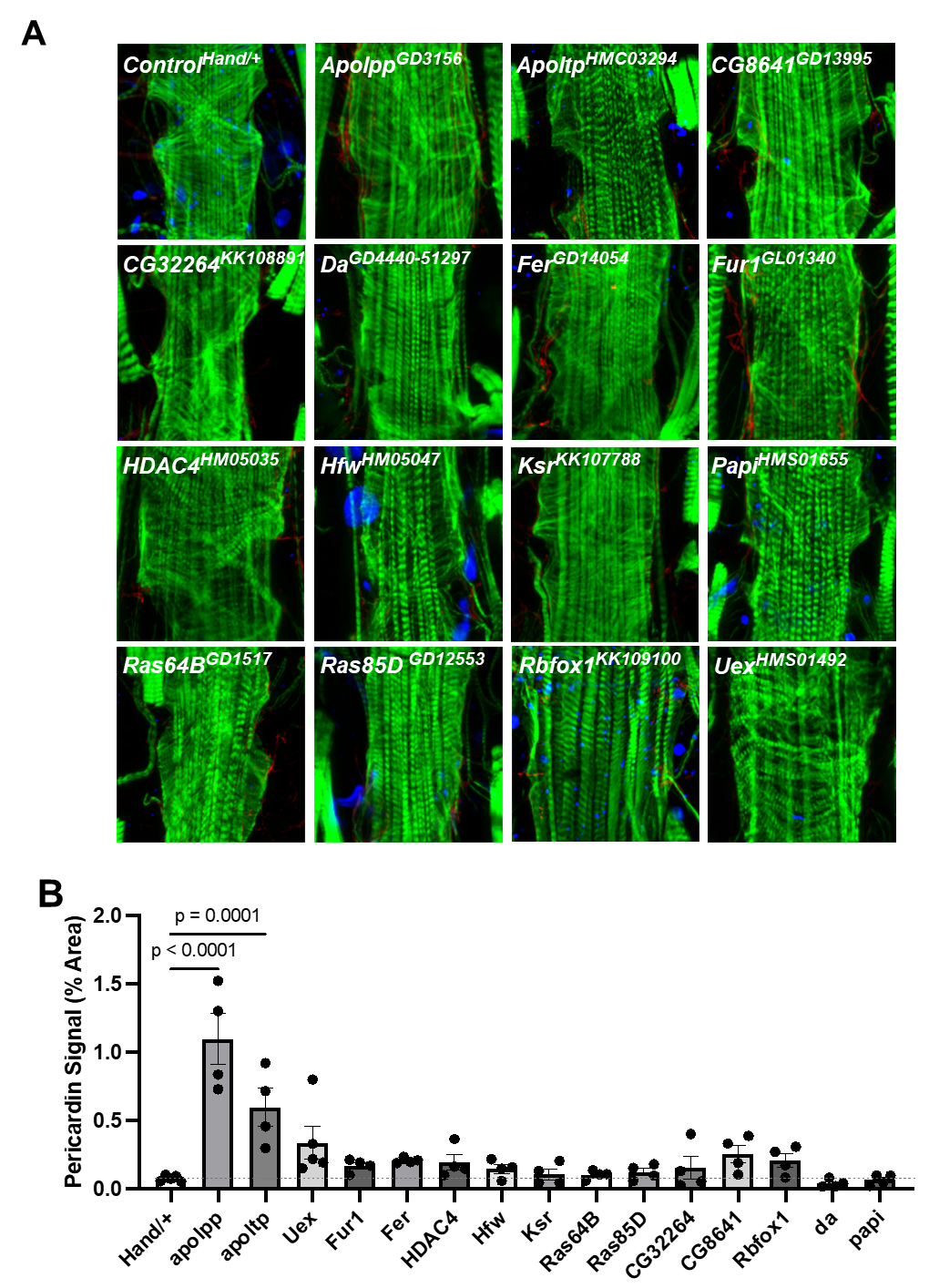

Fig. S3. Cardiac-specific suppression of insomnia- and CVD-related genes leads to myofibrillar disorganization and cardiac fibrosis.

(**A**) Representative images showing actin-containing myofibrils from 3-week-old male flies with cardiac RNAi knockdown of insomnia- and CVD-related genes with *Hand-Gal4*. Each data point is single fly. (**B**) Quantification of pericardin signal. Statistics were calculated by one-way ANOVA with Dunnett’s post hoc test.

Table S1. Regional trait associations for each candidate gene from insomnia- and CVD-associated genetic loci

Regional sleep trait and CVD-related trait associations, in humans, near each candidate gene. CAD (Coronary Artery Disease), DBP (Diastolic Blood Pressure), SBP (Systolic Blood Pressure), HT (Hypertension), and MI (Myocardial Infarction).

Tables S2 – S7 are provided in accompanying .xlsx file

Table S2. Known disease associations for each candidate gene from insomnia- and CVD-associated genetic loci.

Annotation of known disease or syndrome associations, for each candidate genes, with sleep or cardiac features gather from OMIM and additional literature searches.

Table S3. GTEx cis-eQTL associations for each candidate gene from insomnia- and CVD-associated genetic loci.

Significant gene-variant associations indentifed across GTEx. Gencode gene identifier, gene symbol, variant identifier, SNP identifier, association p-value, normalized effect size (NES), and tissue in which the eQTL association was observed are provided for each candidate gene. NES represents the normalized effect size of the alternative allele on gene expression.

Table S4. Orthology of *Drosophila* orthologs for each candidate gene from insomnia- and CVD-associated genetic loci.

Orthologs of each candidate human gene in *Drosophila melanogaster* as obtained from DIOPT. DIOPT orthology scores, orthology rank, human to fly and fly to human orthology from DIOPT version 8 (version at inception of the study) and DIOPT version 10 (current version).

Table S5. UP-TORR data for RNAi lines used in *Drosophila* modeling.

RNAi information for lines used for sleep and heart disruptions in *Drosophila* collected from the Updated Targets of RNAi Reagents (UP-TORR) tool from DRSC (PMID: 23792952). General RNAi information is accompanied by annotation of the background control used as for comparison for each RNAi line, potential off targets, and known insertion site disruptions.

Table S6. Sleep and Cardiac characterization for pan-neuronal knockdown of each *Drosophila* ortholog.

Raw mean and standard deviations for each sleep cand cardiac trait measures along with adjusted Pvalue from one-way ANOVA followed by Šidák’s post hoc test comparing each RNAi to its designated background matched control for *elav-Gal4* mediated knockdowns. N/A or Not Tested indicate an RNAi line that was not tested in *elav-Gal4* experiments due to 40D insertion in a KK line or not prioritized for cardiac phenotyping based on strength of sleep phenotypes present.

Table S7. Sleep and Cardiac characterization for heart knockdown of each *Drosophila* ortholog.

Raw mean and standard deviations for each cardiac and sleep trait measures along with adjusted Pvalue from one-way ANOVA followed by Šidák’s post hoc test comparing each RNAi to its designated background matched control for *Hand-Gal4* mediated knockdowns. N/A or Not Tested indicate an RNAi line that was not prioritized for sleep phenotyping based on strength of cardiac phenotypes present.
